# Two cortical representations of voice control are differentially involved in speech fluency

**DOI:** 10.1101/2020.09.04.283275

**Authors:** Nicole E. Neef, Annika Primaßin, Alexander Wolff von Gudenberg, Peter Dechent, Heiner Christian Riedel, Walter Paulus, Martin Sommer

## Abstract

Recent studies have identified two distinct cortical representations of voice control in humans, the ventral and the dorsal laryngeal motor cortex. Strikingly, while persistent developmental stuttering has been linked to a white matter deficit in the ventral laryngeal motor cortex, intensive fluency shaping intervention modulated the functional connectivity of the dorsal laryngeal motor cortical network. Currently, it is unknown whether the underlying structural network organization of these two laryngeal representations is distinct or differently shaped by stuttering intervention. Using probabilistic diffusion tractography in 22 individuals who stutter and participated in a fluency shaping intervention, in 18 individuals who stutter and did not participate in the intervention, and in 28 control participants, we here compare structural networks of the dorsal laryngeal motor cortex and the ventral laryngeal motor cortex and test intervention-related white matter changes. We show (i) that all participants have weaker ventral laryngeal motor cortex connections compared to the dorsal laryngeal motor cortex network, regardless of speech fluency, (ii) connections of the ventral laryngeal motor cortex were stronger in fluent speakers, (iii) the connectivity profile of the ventral laryngeal motor cortex predicted stuttering severity, (iv) but the ventral laryngeal motor cortex network is resistant to a fluency shaping intervention. Our findings substantiate a weaker structural organization of the ventral laryngeal motor cortical network in developmental stuttering and imply that assisted recovery supports neural compensation rather than normalization. Moreover, the resulting dissociation provides evidence for functionally segregated roles of the ventral laryngeal motor cortical and dorsal laryngeal motor cortical networks.

## 1. Introduction

The human precentral gyrus comprises two representations of voice control. The dorsal laryngeal motor cortex (dLMC) is located between the cortical representations of the lips and the hands (dLMC) (Rödel *et al*., 2004; Brown *et al*., 2008; Olthoff *et al*., 2008; Bouchard *et al*., 2013; Belyk and Brown, 2017). The ventral laryngeal motor cortex (vLMC) occupies parts of the subcentral gyrus and the rolandic operculum (Foerster, 1931; Bouchard *et al*., 2013; Breshears *et al*., 2015). Both regions control articulatory voicing thereby contributing to the production of voiced and voiceless speech sounds, however, the voluntary control of vocal pitch in human speech and singing seems to be selectively encoded via dLMC neurons (Dichter *et al*., 2018). Partly due to technical challenges with investigating speech and singing in vivo in the human brain, only little is known about the structural and functional organization of dLMC and vLMC networks (Simonyan, 2014; Kumar *et al*., 2016; Belyk and Brown, 2017).

Previous *in vivo* imaging studies of the structural connectivity of the laryngeal motor cortex in humans were restricted to the dLMC (Simonyan *et al*., 2009; Kumar *et al*., 2016).

Probabilistic diffusion tractography showed that the human left and right dLMC has moderate connections with the inferior frontal gyrus, superior temporal gyrus, supplementary motor area, caudate nucleus, putamen, and globus pallidus, and particularly dense projections with the somatosensory cortex and the inferior parietal cortex (Kumar *et al*., 2016). All these connections are anatomically plausible and validated by neuroanatomical tract tracing studies of the laryngeal motor cortex representation in the rhesus monkey (Simonyan and Jürgens, 2002, 2003, 2005*a, b*). However, compared to macaque, the human dLMC network showed stronger connections with brain regions involved in the processing of sensory information and feedback, i.e. the primary somatosensory cortex, inferior parietal lobe and superior temporal gyrus (Kumar *et al*., 2016). The authors discuss this finding with the idea that in particular the enhanced connectivity of the dLMC with parietotemporal regions that are involved in sensorimotor integration might have contributed to the development of the sophisticated vocal motor control that is essential for fluent speech production. The dLMC is part of the vast vocal tract sensorimotor cortex and fluent speech production involves the whole orofacial homunculus and in particular, the subcentral gyrus and the rolandic operculum. This ventral extension of the central sulcus harbours the vLMC (Foerster, 1936; Bouchard *et al*., 2013; Breshears *et al*., 2015). Only recently studies start targeting and differentiating findings from the dorsal and the ventral motor representation of voice control (Dichter *et al*., 2018; Belyk *et al*., 2020). Ultimately, fundamental questions exist about what is the structural organization of the vLMC network, does it differ from dLMC network organization, and will the learning of a changed voicing behaviour reorganize the structural network formation within both networks?

One intriguing approach to scrutinize structural network characteristics of the dual cortical laryngeal motor representations is the study of network organization in persistent developmental stuttering. Persistent developmental stuttering is a speech fluency disorder with a complex genetic basis (Kraft and Yairi, 2012). Most often it occurs in early childhood without obvious reason and persists in about 1% of the adults preferably in the male population (Yairi and Ambrose, 2013). Stuttering is evident in sound and syllable repetitions, sound prolongations, and speech blocks, which demonstrates the difficulties of affected individuals to initiate, control and terminate speech movements (Guenther, 2016). These speech motor signs are often accompanied by physical concomitants such as facial grimacing, head and limb movements. Experience of stuttering can cause avoidance behaviours and social anxieties and may impact social well-being, professional career, and socio-economic status (Craig and Tran, 2014). It is widely assumed that stuttering results from a neurofunctional deficit of speech motor planning, sequencing and sensorimotor integration involving system-wide correlates of the speech function, in particular left perisylvian speech areas, basal ganglia, and cerebellum (Ludlow and Loucks, 2003; Craig-McQuaide *et al*., 2014; Neef *et al*., 2015; Etchell *et al*., 2017; Chang *et al*., 2018; Connally *et al*., 2018; Chang and Guenther, 2019). Strikingly, one robust neural trait marker of persistent developmental stuttering is a white matter deficit adjacent to the left vLMC (Sommer *et al*., 2002; Watkins *et al*., 2008; Chang *et al*., 2010; Neef *et al*., 2015). Involved fiber tracts connect fronto-parietal/temporal circuit that promote speech production (Hickok and Poeppel, 2007; Friederici, 2011; Hickok, 2012). A disruption of these connections might disturb speech signal transmission and thus, hamper speech fluency (Sommer *et al*., 2002). On the contrary to the white matter deficit in the left vLMC, fluency shaping, a stuttering intervention that involves learning to speak with reduced pitch modulation and voicing complexity (Euler *et al*., 2009), synchronizes task-free brain activity between the left dLMC and sensorimotor brain regions (Korzeczek *et al*., 2020). Briefly summarized, persistent stuttering is linked to a white matter deficit in the left vLMC, while assisted recovery from stuttering via fluency shaping is linked to an increased functional connectivity of the dLMC.

Currently, it is an open question, whether intensive learning of a new voicing pattern will shape the structural organisation of the two laryngeal motor representations and if so whether neuroplasticity is similar or different between these two networks. In addition, it is in general unclear, whether dLMC and vLMC have structural connectivity patterns that are distinct or comparable, independent from the speech fluency of studied individuals. Therefore, in the present study, we re-examined diffusion MRI data of healthy adult humans, adults who stutter and adults who stutter and who participated in an 11-month intensive fluency-shaping intervention. We used probabilistic diffusion tracking with bilateral seeds in the dLMC and vLMC and quantified respective connection probabilities with the somatosensory cortex, inferior parietal cortex, inferior frontal gyrus, superior temporal gyrus, supplementary motor area, caudate nucleus, putamen, and globus pallidus (Kumar *et al*., 2016). Neuroanatomical tract tracing studies of the laryngeal motor cortex representation in the rhesus monkey show in addition hard-wired reciprocal connections with the thalamus, anterior cingulate cortex and midcingulate cortex (Jürgens, 2002; Simonyan *et al*., 2009; Price, 2012; Simonyan and Fuertinger, 2015). However, in a previous study these target regions revealed no significant proportions of projections when applying probabilistic diffusion tracking (Kumar *et al*., 2016) and thus were not included in the current analyses. We used an exploratory statistical approach, i.e. mixed-model ANCOVA and required downstream post-hoc statistics, to determine the influence of seed, hemisphere, target, time and group. Furthermore, we tested whether stuttering severity was predicted by the structural network profiles of dLMC and vLMC, respectively.

## 2. Materials and methods

### 2.1 Participants

Current data were derived from a dissertation project (Primaßin, 2019) that evaluated the long-term effects of an intensive stuttering intervention on white matte integrity and task-related brain activity. Here, we analyzed diffusion MRI data sets of 22 adults with stuttering who took part in a fluency-shaping program (AWS+, 2 females, mean age 25.6 ± 11.7 SD), 18 adults with stuttering who did not participate in any intervention during this study (AWS-, 2 females, mean age 34.8 ± 7.0 SD), and 28 adults without stuttering (AWNS, 4 females, mean age 25.1 ± 7.4 SD). Participants completed two MRI sessions 11.5 ± 1.1 SD month apart and received an allowance for their expenses. All were monolingual native speakers of German, reported normal (or-corrected-to normal) vision and no history of hearing, speech, language or neurological deficits apart from stuttering in the AWS groups, drug abuse, or medications that act on the central nervous system. The groups were matched for sex and handedness(Oldfield, 1971). AWS-were older and had a higher education score than participants in the two other groups (see Table 1). Education and age were correlated with *r* = 0.483, *p* < 0.001, and therefor only age was considered as a covariate in all statistical analyses.

**Table 1.**
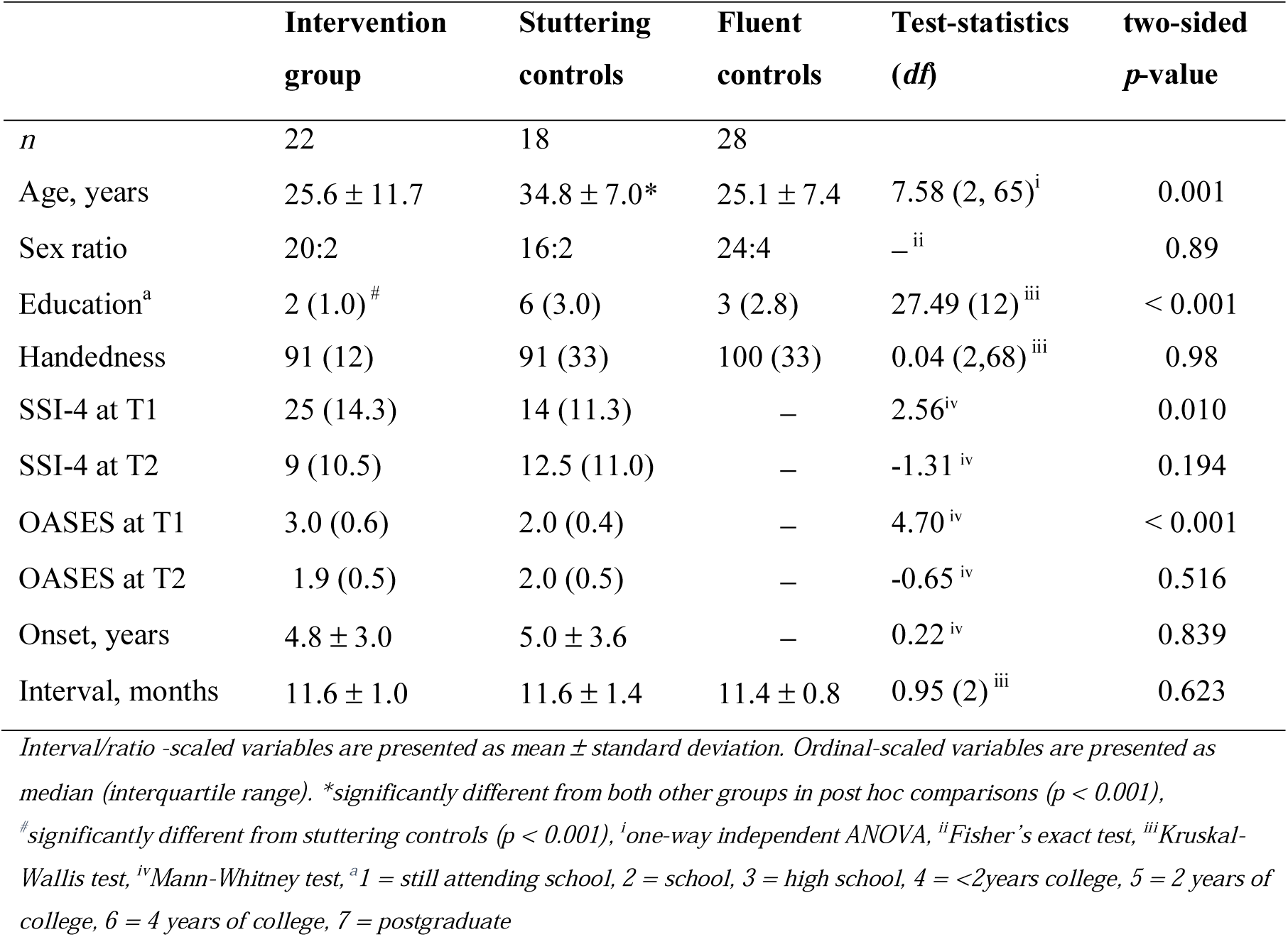
Demographic information of participants

|  | **Intervention group** | **Stuttering controls** | **Fluent controls** | **Test-statistics (df)** | **two-sided p-value** |
| --- | --- | --- | --- | --- | --- |
| *n* | 22 | 18 | 28 |  |  |
| Age, years | 25.6 ± 11.7 | 34.8 ± 7.0* | 25.1 ± 7.4 | 7.58 (2, 65)<sup>i</sup> | 0.001 |
| Sex ratio | 20:2 | 16:2 | 24:4 | —<sup>ii</sup> | 0.89 |
| Education<sup>a</sup> | 2 (1.0)<sup>#</sup> | 6 (3.0) | 3 (2.8) | 27.49 (12)<sup>iii</sup> | < 0.001 |
| Handedness | 91 (12) | 91 (33) | 100 (33) | 0.04 (2,68)<sup>iii</sup> | 0.98 |
| SSI-4 at T1 | 25 (14.3) | 14 (11.3) | — | 2.56<sup>iv</sup> | 0.010 |
| SSI-4 at T2 | 9 (10.5) | 12.5 (11.0) | — | -1.31<sup>iv</sup> | 0.194 |
| OASES at T1 | 3.0 (0.6) | 2.0 (0.4) | — | 4.70<sup>iv</sup> | < 0.001 |
| OASES at T2 | 1.9 (0.5) | 2.0 (0.5) | — | -0.65<sup>iv</sup> | 0.516 |
| Onset, years | 4.8 ± 3.0 | 5.0 ± 3.6 | — | 0.22<sup>iv</sup> | 0.839 |
| Interval, months | 11.6 ± 1.0 | 11.6 ± 1.4 | 11.4 ± 0.8 | 0.95 (2)<sup>iii</sup> | 0.623 |

*Interval/ratio -scaled variables are presented as mean ± standard deviation. Ordinal-scaled variables are presented as median (interquartile range). \*significantly different from both other groups in post hoc comparisons ( $p < 0.001$ ), <sup>#</sup>significantly different from stuttering controls ( $p < 0.001$ ), <sup>i</sup>one-way independent ANOVA, <sup>ii</sup>Fisher's exact test, <sup>iii</sup>Kruskal-Wallis test, <sup>iv</sup>Mann-Whitney test, <sup>a</sup>1 = still attending school, 2 = school, 3 = high school, 4 = <2years college, 5 = 2 years of college, 6 = 4 years of college, 7 = postgraduate*

The ethical review board of the University Medical Center Göttingen, Georg August University Göttingen, Germany, approved the study, and all participants provided written informed consent, according to the Declaration of Helsinki, before participation.

Speech fluency of all participants, determined by using the Stuttering Severity Index (SSI-4, (Riley *et al*., 2004), was assessed prior to each MRI session. As part of this assessment, each AWS was video recorded while reading aloud and speaking with an experimenter. Two certified speech-language pathologist (one of them was A.P.) then rated the frequency and durations of the stuttered syllables and the presence of physical concomitants. At test time point one (T1), stuttering severity in the AWS+ group ranged from 7 to 39, with a median of 25 and an interquartile range of 15 to 31. Five of the 22 AWS+ were categorized as very mild, 5 as mild, 6 as moderate, 3 as severe, 2 as very severe, and one with an SSI-4 total score of 7 was not classified. The stuttering severity in the AWS-group ranged from 4 to 42, with a median of 14 and an interquartile range of 7 to 21. Eight of the 18 AWS-were categorized as very mild, 2 as mild, 1 as moderate, 1 as severe, 1 as very severe, and 5 with SSI-4 scores between 4 and 7 were not classified. Fluency-shaping reduced stuttering severity in AWS+,(Primaßin, 2019; Korzeczek *et al*., 2020). Accordingly, at test time point two (T2), stuttering severity in the AWS+ group ranged from 1 to 37, with a median of 9 and an interquartile range of 5 to 16. After therapy 5 of the AWS+ group were categorized as very mild, 2 as mild, 1 as moderate, 1 as very severe, and 13 with an SSI-4 score smaller than 9 were not classified. Before intervention, stuttering severity was more severe in AWS+ as compared to AWS-. Similarly, the self-assessment of the psycho-social impact of stuttering (Overall Assessment of the Speaker’s Experience of Stuttering, OASES)(Yaruss and Quesal, 2014) indicated that the participants of the intervention group suffered more from stuttering than stuttering controls. These group differences vanished after stuttering intervention (see Table 1).

### 2.2 Image Acquisition

MRI data were acquired in a 3 Tesla Siemens Magnetom Tim Trio scanner (Erlangen, Germany) using an eight-channel phased-array head coil at the University Medical Center Göttingen, Germany. Sagittal T1 weighted structural data were acquired with a 3D turbo fast low angle shot (FLASH) sequence (TR = 2250ms, TE = 3.26ms, TI = 900ms, flip angle = 9°, 256mm FoV, 7/8 Fourier phase encoding) as whole-brain anatomical reference data at a spatial resolution of 1 × 1 × 1 mm^3^ voxel size (256 × 256 matrix). Diffusion-weighted MRI data were acquired with a spin-echo EPI sequence (TR = 10100 ms, TE = 93 ms, parallel acquisition factor 2, 6/8 Fourier phase encoding, 243mm FoV, acquisition matrix: 128 × 128, 74 slices, voxel size 1.9 × 1.9 × 1.9 mm^3^) sampling 64 image volumes with diffusion weighting along 64 diffusion directions (b = 1000 s/mm^2^) and one reference image without diffusion weighting. Participants lay in supine position in the scanner and wore headphones for noise protection, and MR-compatible LCD goggles (VisuaStim XGA, Resonance Technology Inc., Northridge, CA, USA).

### 2.3 MRI Data Analysis

Diffusion-weighted (d)MRI images were processed with FSL, http://www.fmrbi.ox.ac.uk/fsl/(Jenkinson *et al*., 2012). Images were corrected for eddy currents and head motion by using affine registration to the non-diffusion volumes.

Probabilistic tractography was performed in the native dMRI space. We computed voxel-wise estimates of the fiber orientation distribution of up to two fiber orientations with the FSL function bedpost.(Behrens *et al*., 2007; Jbabdi *et al*., 2012) Seed and target masks were 3 mm spheres. Coordinates of seeds of the dLMC were derived from a previous quantitative meta-analysis (Kumar *et al*., 2016) but shifted from the gray matter in the anterior wall of the central sulcus [x = -45, y = -14, z = 33; x = 44, y = -12, z = 35] to the white matter in the precentral gyrus [x = -47, y = -4, z = 34; x = 45, y = -3, z = 35] according to the FSL_HCP1065_FA _1mm standard image. Seeds of the vLMC were placed at [x = -46, y = - 16, z = 19; x = 46, y = -16, z = 19] in the white matter adjacent to the subcentral sulcus and the rolandic operculum (Sommer *et al*., 2002). Target masks of the supplementary motor area (SMA), inferior frontal gyrus pars opercularis (IFGop), inferior frontal gyrus pars triangularis (IFGtr), primary somatosensory cortex (S1), inferior parietal lobe (IPL), anterior superior temporal gyrus (aSTG), putamen (Put), caudate nucleus (Caud), and globus pallidus (Gp), listed in Table 2, were placed at the center of gravity of maximal tract probability derived from a probabilistic diffusion tractography in humans (Kumar *et al*., 2016), and warped to the native dMRI (Andersson *et al*., 2010). We used modified Euler streamlining, distance correction, and 100,000 samples per voxel within the FSL function probtrackx2 with three pairs of seed and target mask. Target mask determined both waypoint and termination mask to compute the structural connectivity between left hemispheric brain regions. All analyses were calculated separately for each pair of seed and target region. The ‘connectivity index’ was determined from the number of sample streamlines from each seed that reached the target. We normalized the connectivity index by dividing the logarithm of the number of streamlines from a given seed that reached the target (i.e., numeric output of the tractography algorithm given as waytotal) by the logarithm of the product of the number of generated sample streamlines in each seed voxel (100,000) and the number of voxels in the seed mask, n = 19. The logarithmic scaling transformed the connectivity index into a normally distributed variable with a range between 0 and 1.

**Table 2.**
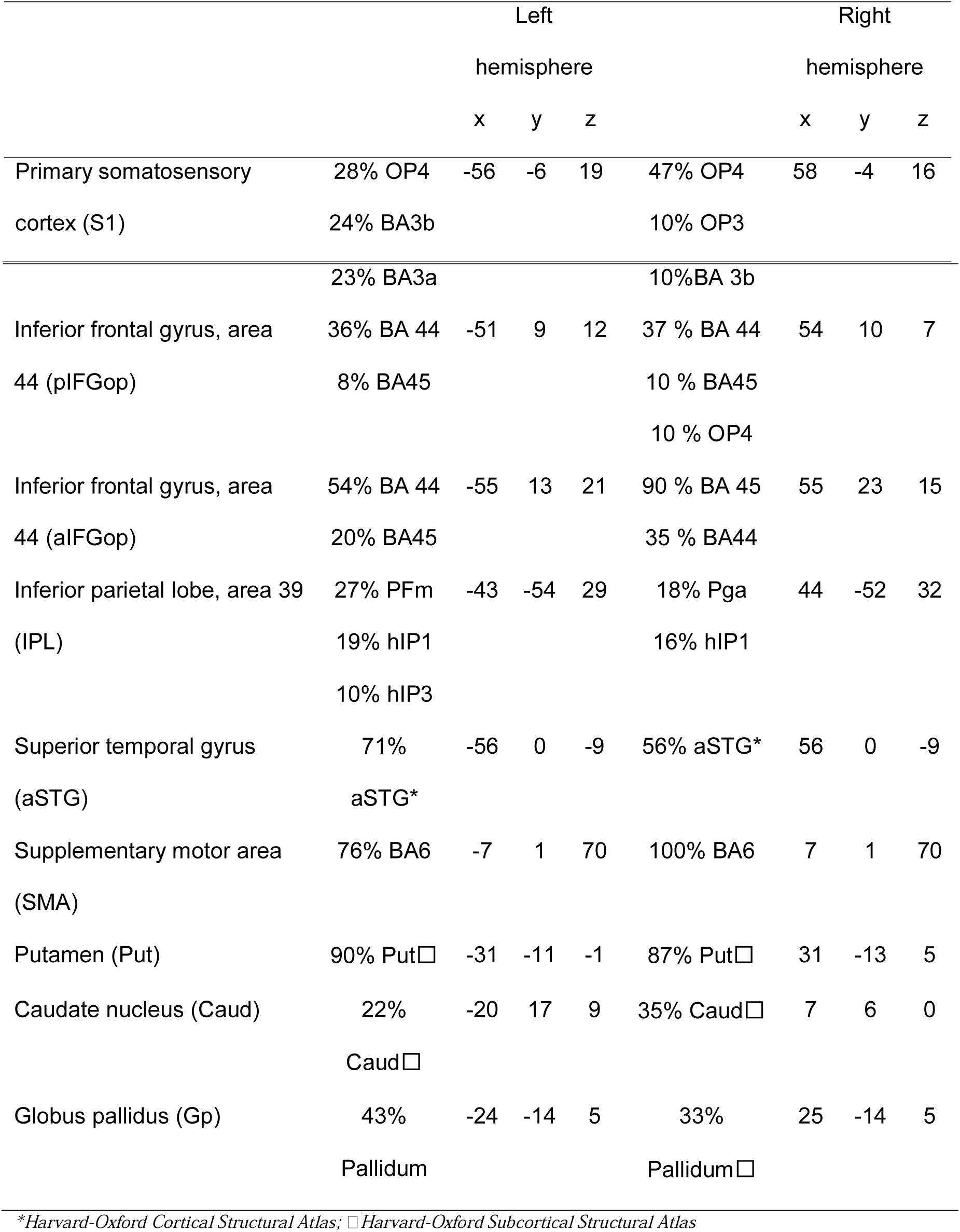
MNI coordinates of target regions

### 2.4 Statistical analyses

For quantitative between-group analysis of the human dLMC and vLMC network we used the connectivity indices revealed for each seed-to-target pair within one mixed model ANCOVA. We modeled Group (AWS+, AWS-, AMNS) as between-subjects factor, Time (T1, T2), Seed (dLMv, vLMV), Hemisphere (left hemisphere, right hemisphere), and Target region (SMA, pIFGop, aIFGop, S1, IPL, STG, Put, Caud, Gp) as repeated measures within-subjects factors, and Age as a covariate. If the main effect of Seed was significant, the follow-up post hoc ANCOVAs examined the two LMC networks separately with Group (AWS+, AWS-, AMNS) as between-subjects factor, Time (T1, T2), Hemisphere (left hemisphere, right hemisphere), and Target region (SMA, aIFGop, pIFGop, S1, IPL, STG, Put, Caud, Gp) as repeated measures within-subjects factors, and Age as a covariate.

We assessed tract lateralization by using the laterality index calculated as (connectivity index in the right hemisphere – connectivity index in the left hemisphere)/(connectivity index in the right hemisphere + connectivity index in the left hemisphere). A positive value would indicate a lateralization to the right, whereas a negative value would indicate a lateralization to the left. We tested the significance of the lateralization by calculating two-sided paired *t*-tests against a mean value of zero and report significant lateralization at p < 0.05 (Bonferroni-corrected). Furthermore, we compared laterality indices between seeds with paired *t*-tests and report significant differences at p < 0.05 (Bonferroni-corrected).

Hierarchical regressions were used to predict pre-intervention speech fluency of affected individuals from structural connectivity profiles of dLMC and vLMC, respectively. The first step comprised demographic factors (age, sex, and handedness) and the second step comprised connection probabilities; thus, the models estimate what percentage of variance in structural connectivity accounts for speech fluency above and beyond demographics.

### 2.5. Data availability statement

The data that support the findings of this study are available from the corresponding author upon reasonable request.

## 3. Results

### 3.1 GLM

The 3 × 2 × 2 × 2 × 9 ANCOVA performed on the connection strength of dLMC and vLMC revealed a significant main effect of seed, *F*(1,64) = 11.821, *p* = .001, *η*^2^_*p*_ =.156, such that dLMC, mean (*M*) = .377, confidence interval 95% CI [.367 - .388] is characterized with an overall higher connectivity than vLMC, *M* = .315, 95% CI [.306 – .325]. There was also a trend for an interaction of Seed × Group, *F*(2,64) = 2.728, *p* = .0073, *η*^2^_*p*_ =.079. In addition, there were a main effect of Target region, a main effect of Age and an interaction of Seed × Hemisphere, an interaction of Seed × Target region, an interaction of Hemisphere × Target region, an interaction of Hemisphere × Target region × Age, an interaction of Seed × Hemisphere × Target region, and an interaction of Seed × Hemisphere × Target region × Age, all reported in Table 3. The 4-way interaction indicates that the structural connectivity of dLMC and vLMC vary depending on hemisphere and target region and that this variance is in addition modulated by age. There was no main effect of Time or interaction of Time × Seed × Group or of Time × Seed × Target region × Group indicating no change of the structural connectivity of the two larynx areas over time and no intervention-induced neuroplasticity at this global analysis level.

**Table 3.** Results of the global mixed model ANCOVA

| | *df* | *F* | *p* | $\eta^2_p$ |
| --- | --- | --- | --- | --- |
| Seed | 1 | 11.821 | 0.001 | 0.156 |
| Seed × Group | 2 | 2.728 | 0.073 | 0.079 |
| Target region | 6.215 | 33.157 | < 0.001 | 0.341 |
| Seed × Hemisphere | 1 | 2.967 | 0.090 | 0.044 |
| Seed × Target region | 5.845 | 20.253 | < 0.001 | 0.240 |
| Hemisphere × Target region | 5.078 | 3.703 | 0.003 | 0.055 |
| Hemisphere × Target region × Age | 8 | 2.067 | 0.037 | 0.031 |
| Seed × Hemisphere × Target region | 5.344 | 2.868 | 0.013 | 0.043 |
| Seed × Hemisphere × Target region × Age | 8 | 2.785 | 0.015 | 0.042 |
| Age | 1 | 9.433 | 0.003 | 0.128 |

**Table 4.**
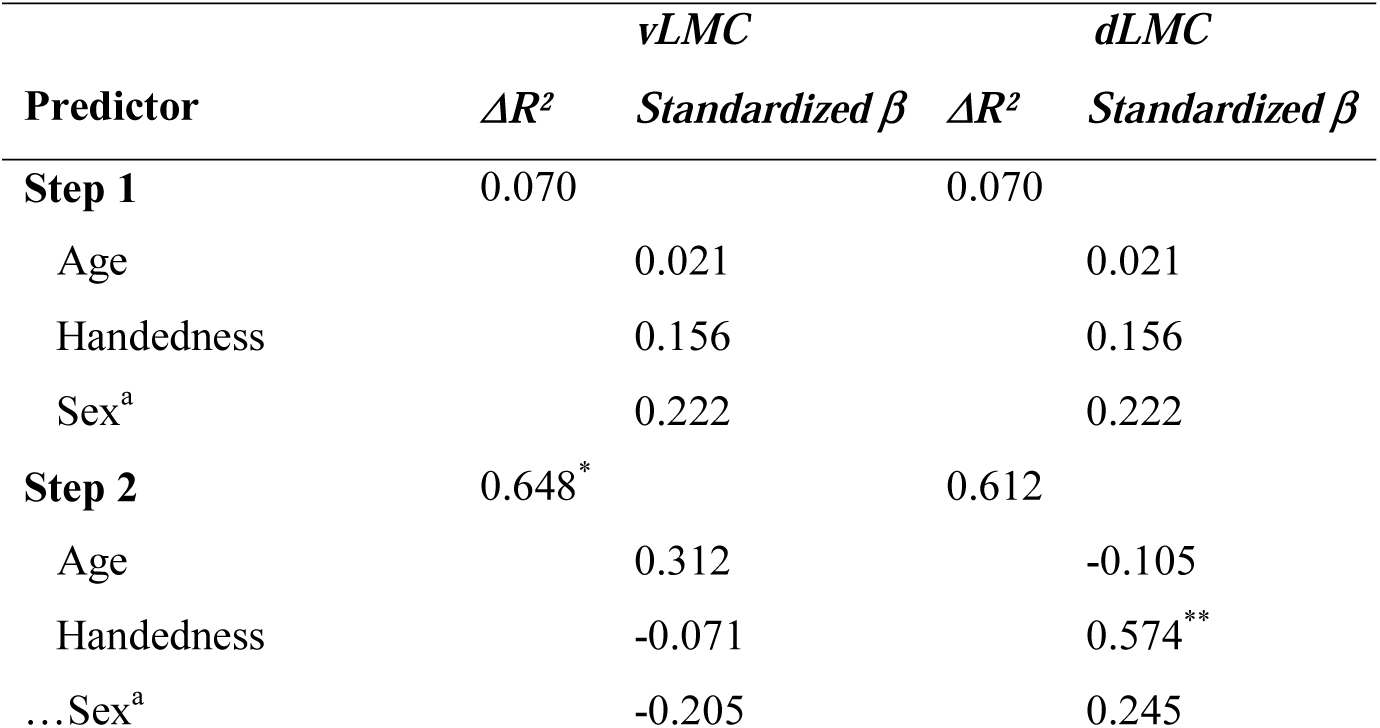
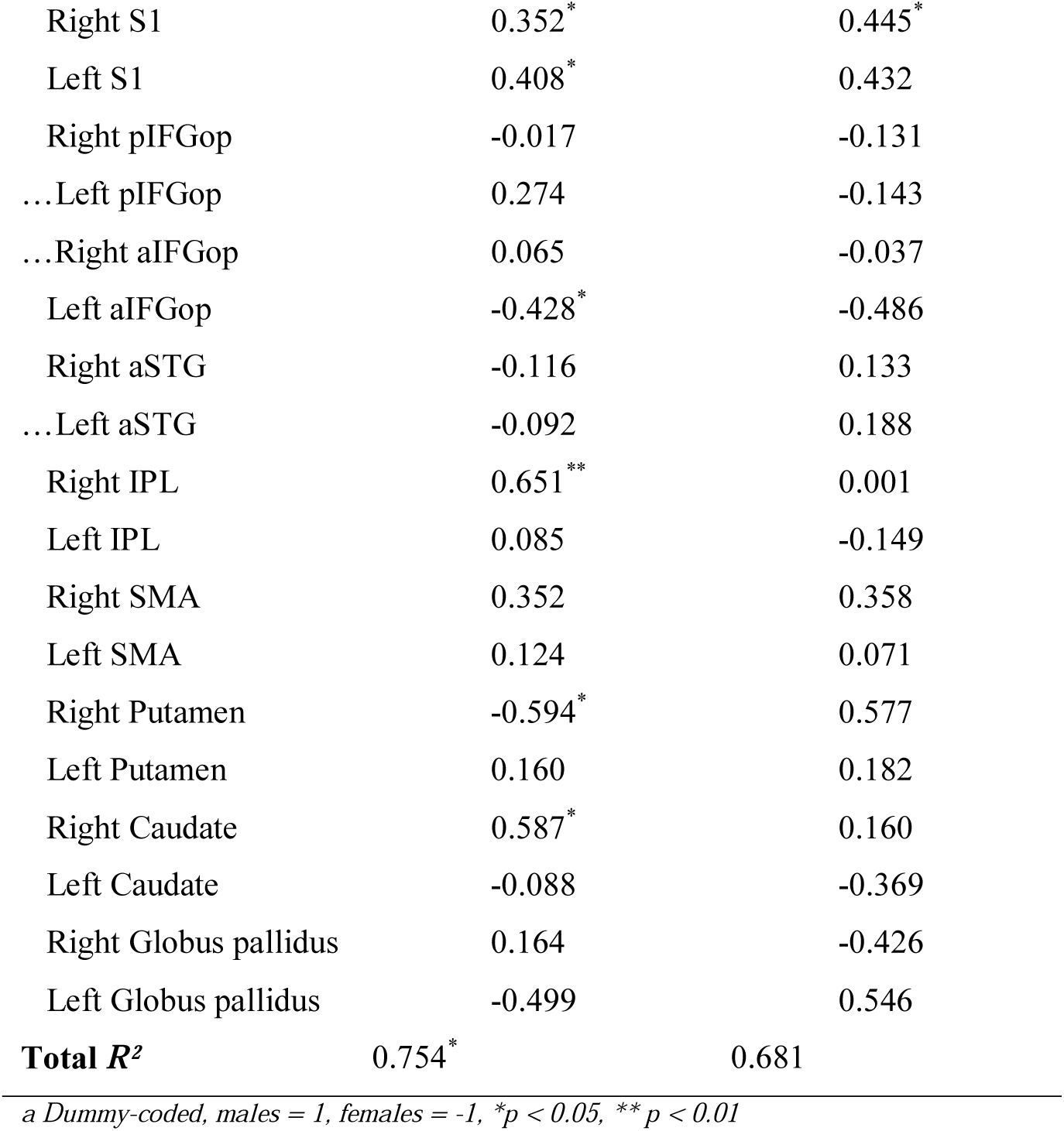
Connection probability of vLMC predicts motor signs of stuttering

| Predictor | vLMC |  | dLMC |  |
| --- | --- | --- | --- | --- |
| | $\Delta R^2$ | Standardized $\beta$ | $\Delta R^2$ | Standardized $\beta$ |
| **Step 1** | 0.070 |  | 0.070 |  |
| Age |  | 0.021 |  | 0.021 |
| Handedness |  | 0.156 |  | 0.156 |
| Sex<sup>a</sup> |  | 0.222 |  | 0.222 |
| **Step 2** | 0.648<sup>*</sup> |  | 0.612 |  |
| Age |  | 0.312 |  | -0.105 |
| Handedness |  | -0.071 |  | 0.574<sup>**</sup> |
| ...Sex<sup>a</sup> |  | -0.205 |  | 0.245 |

|  |  |  |
| --- | --- | --- |
| Right S1 | 0.352<sup>*</sup> | 0.445<sup>*</sup> |
| Left S1 | 0.408<sup>*</sup> | 0.432 |
| Right pIFGop | -0.017 | -0.131 |
| ...Left pIFGop | 0.274 | -0.143 |
| ...Right aIFGop | 0.065 | -0.037 |
| Left aIFGop | -0.428<sup>*</sup> | -0.486 |
| Right aSTG | -0.116 | 0.133 |
| ...Left aSTG | -0.092 | 0.188 |
| Right IPL | 0.651<sup>**</sup> | 0.001 |
| Left IPL | 0.085 | -0.149 |
| Right SMA | 0.352 | 0.358 |
| Left SMA | 0.124 | 0.071 |
| Right Putamen | -0.594<sup>*</sup> | 0.577 |
| Left Putamen | 0.160 | 0.182 |
| Right Caudate | 0.587<sup>*</sup> | 0.160 |
| Left Caudate | -0.088 | -0.369 |
| Right Globus pallidus | 0.164 | -0.426 |
| Left Globus pallidus | -0.499 | 0.546 |
| **Total R<sup>2</sup>** | **0.754<sup>\*</sup>** | **0.681** |

*a Dummy-coded, males = 1, females = -1, \* $p < 0.05$ , \*\* $p < 0.01$*

### 3.2 Lateralization

Paired *t*-tests assessing the lateralisation of dLMC target regions found aSTG (*t* = 5.59, *p* < 0.001), IPL (*t* = 5.80, *p* < 0.001) and Put (*t* = 3.94, *p* = 0.002) to show greater right-hemispheric connectivity and S1 (*t* = -5.31, *p* < 0.001), aIFGop (*t* = -2.88, *p* < 0.048) and Caud (*t* = -4.64, *p* < 0.001) to show greater left-hemispheric connectivity. Paired *t*-tests assessing the lateralisation of vLMC target regions found aSTG (*t* = 8.76, *p* < 0.001), S1 (*t* = 4.45, *p* < 0.001) and pIFGop (*t* = 4.55, *p* < 0.001) to show greater right-hemispheric connectivity and Caud (*t* = -4.05, *p* < 0.001) to show greater left-hemispheric connectivity. Figure 2 shows LIs separated for seed and target regions.

**Figure 1.**
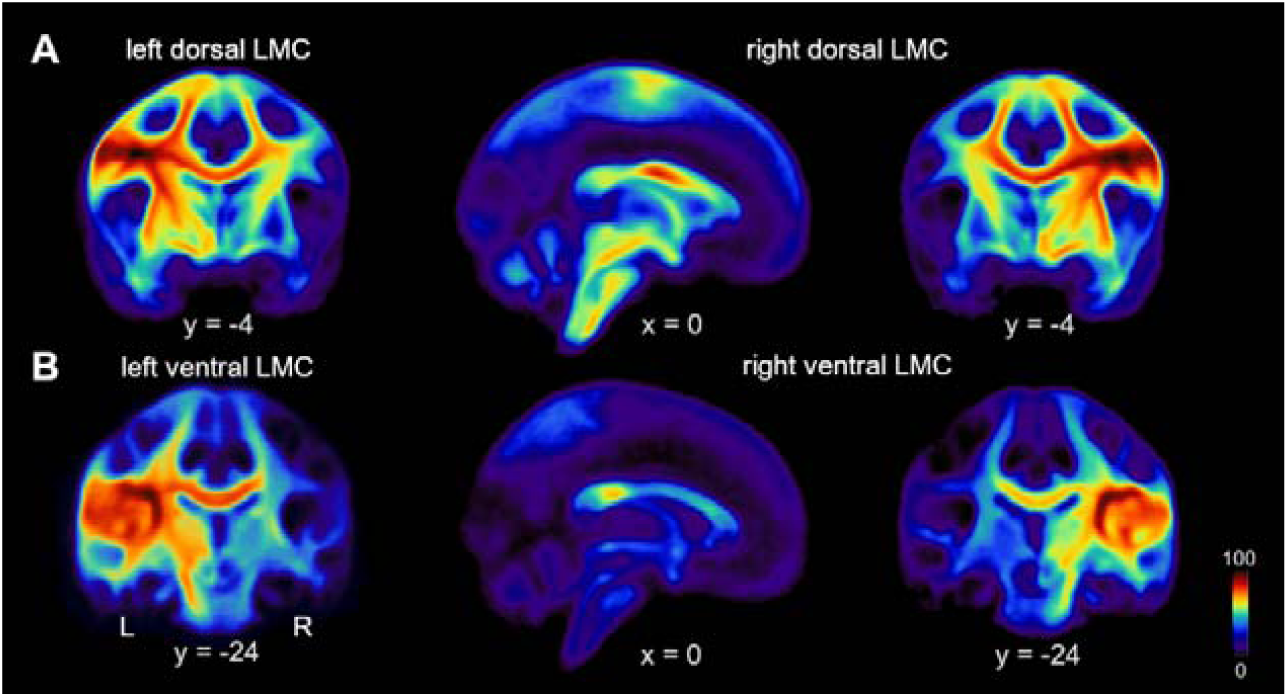
Connection probability of two distinct larynx cortical representations. A) Population probability maps illustrating the likelihood of structural connectivity of the left and right dorsal laryngeal motor cortex (dLMC) and B) left and right ventral laryngeal motor cortex (vLMC) with dark-red marking 100% and dark-blue 0% connection probabilities.

**Figure 2.**
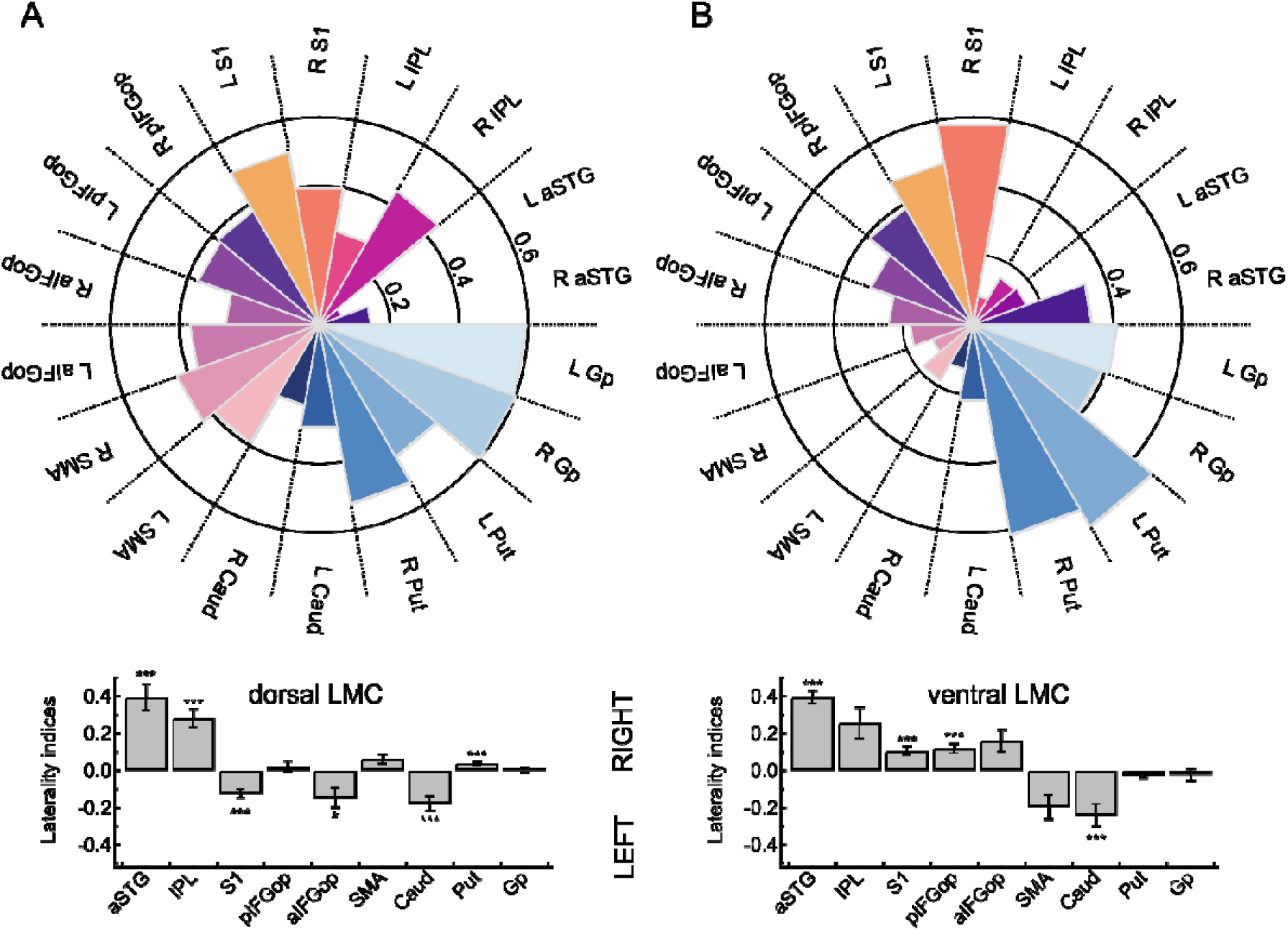
Connection probability fingerprints and hemispheric lateralization of two laryngeal motor representations. (A) Connectivity fingerprints show the likelihood (0-1) of the dorsal laryngeal motor cortex (dLMC) and the (B) ventral laryngeal motro cortex (vLMC) averaged per target region across all participants and all sessions. Bar plots indicate hemispheric lateralization at ***p < 0.001 and * p < 0.05 (Bonferroni-corrcted). Abbreviations: aSTG anterior superior temporal gyrus, Caud = nucleus caudatus, Gp = globus pallidus, pIFGop = posterior inferior frontal gyrus pars opercularis, aIFGop = anterior inferior frontal gyrus pars opercularis, IPL = inferior parietal lobule, Put = putamen, S1 = somatosensory cortex, SMA = supplementary motor area.

Paired *t*-tests assessing whether lateralization differed between dLMC and vLMC were significant for S1 (*t* = -6.90, *p* < 0.001), IFGtr (*t* = -3.89, *p* = 0.001), SMA (*t* = 3.70, *p* = 0.003) and Put (*t* = 4.22, *p* < 0.001) and marginal significant for pIFGop (*t* =-2.72, *p* = 0.067).

### 3.3 Seed-wise GLM Analyses

Because the global ANCOVA revealed a main effect of Seed, we examined the connection strength of the two seed regions separately. The 3 × 2 × 2 × 9 ANCOVA performed on the connection indices of vLMC revealed a main effect of group with *F*(2,64) = 4.843, *p* = .011 such that vLMC connectivity in AWNS, mean (*M*) = .336, confidence interval 95% CI [.321 - .350] was higher than in AWS+, *M* = .309, 95% CI [.293 – .326], and AWS-, *M* = .301, 95% CI [.282 – .321], see Figure 4C. Post-hoc group comparisons revealed significant differences between AWNS and AWS+ with a mean difference of .035 ± .130 SEM, *p* = .008, and AWNS and AWS-with a mean difference of .027 ± .011, *p* = .018, but no difference between AWS+ and AWS-with a mean difference of -.008 ± .013, *p* = .545. Figure 3 illustrates the vLMC connectivity fingerprints for AWS and ANS. There was also a main effect of Target region with *F*(8,64) = 41.738, *p* < .001, *η*^2^_*p*_ =.395, a main effect of age with *F*(1,64) = 6.500, *p* = .013, *η*^2^_*p*_ =.092,, *η*^2^_*p*_ =.131, an interaction of Hemisphere × Target region × Age with *F*(8,64) = 3.672, *p* < .001, *η*^2^_*p*_ =.054, and a trending interaction of Hemisphere × Target region with *F*(4.513,64) = 2.239, *p* = .057, *η*^2^_*p*_ =.034. All other main effects and interactions were not significant [p > .1, in all cases].

**Figure 3.**
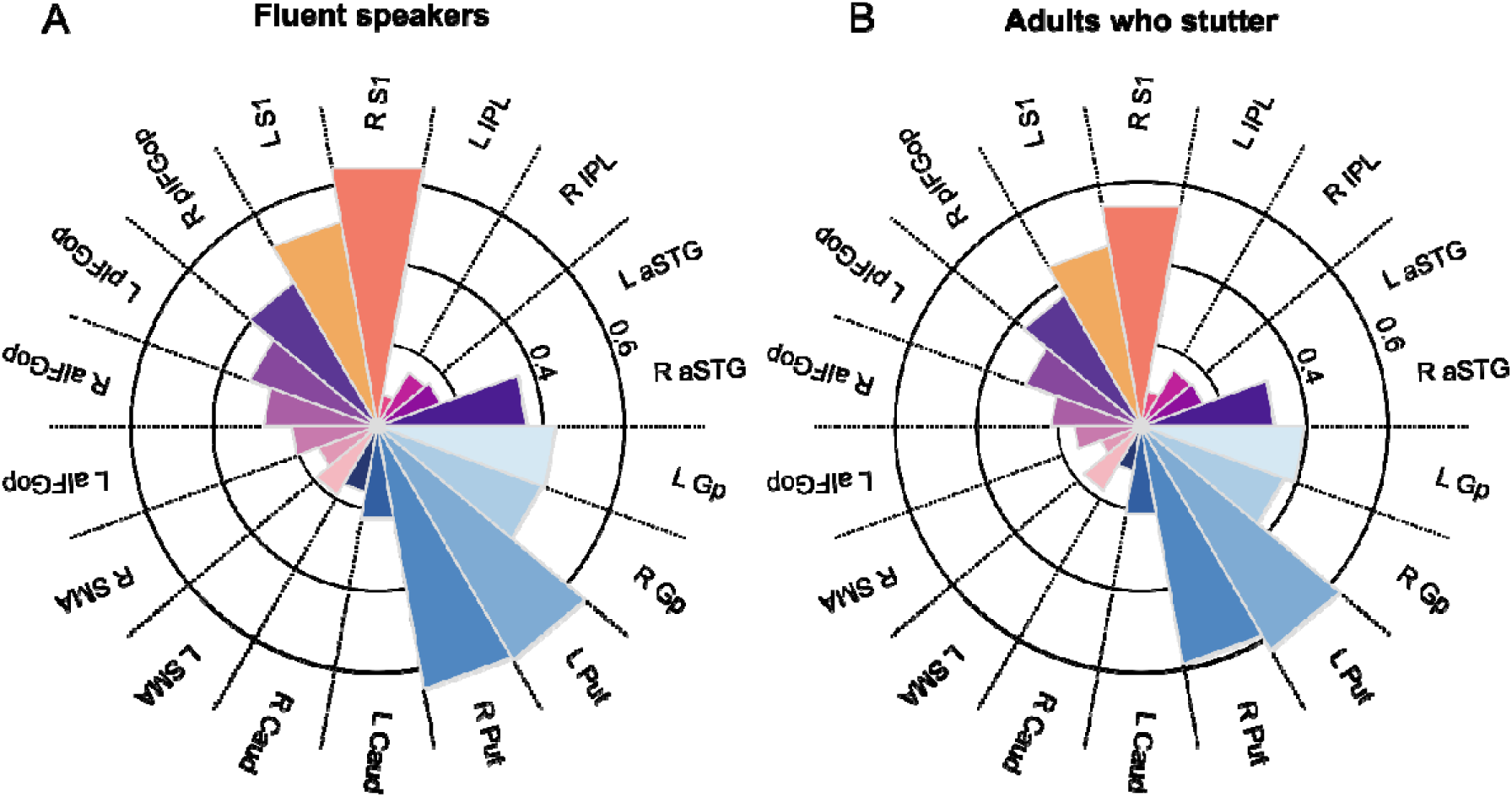
Connection probability fingerprints of the ventral laryngeal motor cortex (vLMC) (A) for fluent speakers and (B) adults who stutter.

**Figure 4.**
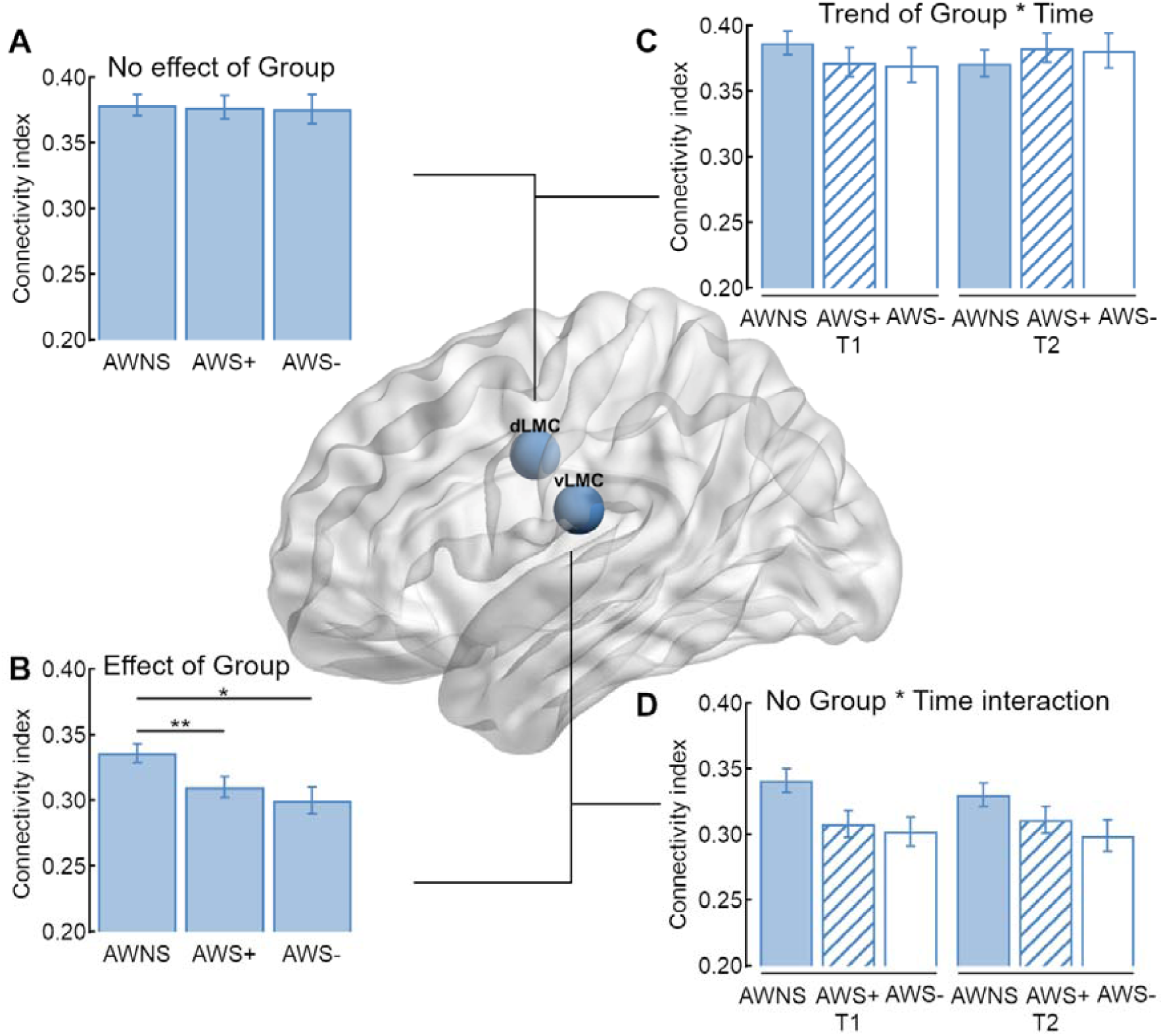
Structural connectivity of the two laryngeal motor cortices. (A) In sum, adults who stutter (AWS) and adults who do not stutter (AWNS) have an overall comparable connectivity index of the dLMC network, (B) but AWS have a decreased overall connectivity index of the vLMC network compared to AWNS. (C) The trending interaction of Group × Time was not driven by an increase of the overall connectivity index of AWS with intensive stuttering intervention (AWS+), but by a trending decrease in AWNS. (D) Time had no influence on the overall structural connectivity of the vLMC network.

The 3 × 2 × 2 × 9 ANCOVA performed on the connection strength of dLMC revealed a main effect of Target region with *F*(5.676,64) = 18.429, *p* < .001, *η*^2^_*p*_ =.224, an interaction of Hemisphere × Target region with *F*(5.152,64) = 4.568, *p* < .001, *η*^2^_*p*_ =.067, a main effect of age with *F*(1,64) = 6.293, *p* = .01, *η*^2^_*p*_ = .090, and a trend towards an interaction of Time × Group with *F*(2,64) = 2.654, *p* = .078, *η*^2^_*p*_ = .077. All other main effects and interactions were not significant [p > .1, in all cases]. To test the interaction of Group × Time we calculated further post-hoc ANCOVAs. For AWNS the analysis revealed a trending effect of time with *F*(1,24) = 3.257, *p* = .082, *η*^2^_*p*_ = .119, but no significant effect or trend for the other two groups. Figure 4C shows a trend towards a decreased overall structural connectivity in AWNS.

### 3.4 Regression analyses

We constructed two statistical models incorporating the connection probability of vLMC and dLMC, respectively, in this cohort of 40 adults with chronic persistent stuttering since childhood. We found that connection probability of the vLMC predicted motor signs of stuttering severity, as measured with the SSI-4, over and above the biological factors age, sex and handedness. Variance of SSI-4 total scores in the cohort was explained with *ΔR2* = 0.648, *F*[18,36] = 2.625, *p* = 0.022; total *R2* = 0.754, *F*[21,39] = 2.779, *p* = 0.018. Contrastingly, connection probability of the dLMC did not predict stuttering severity over and above the biological variates *ΔR2* = 0.612, *F*[18,36] = 1.834, *p* = 0.099; total *R2* = 0.681, *F*[21,39] = 1.920, *p* = 0.088.

## 4. Discussion

One essential component of natural fluent speech is the flexible control of pitch and voicing. This speech function is distributed to two laryngeal representations per hemisphere, the dLMC and vLMC. Here, we show that (1) these cortical representations diverge in their structural connectivity profiles, (2) the dLMC network shares denser connections compared to the vLMC network, (3) the vLMC connectivity is stronger in fluent speakers compared to adults who stutter, (4) the connectivity profile of the vLMC predicts stuttering severity, and (5) none of the two structural LMC networks changed with fluency shaping, a common stuttering intervention with a remarkable change of voice control during speaking.

Our findings indicate that the dLMC has an overall stronger structural connectivity compared to the vLMC. This is in line with a neuroimaging study characterizing the cortical microstructure underlying the two laryngeal representations with quantitative MRI (Eichert *et al*., 2020). Multiparameter mapping and myelin mapping revealed that the dLMC has a myelin content and a cortical thickness that equals that of the primary motor cortex (Fischl and Dale, 2000; Glasser and Van Essen, 2011). Furthermore, myelin content and cortical thickness of the dLMC was higher compared to the vLMC (Eichert *et al*., 2020). The authors discuss their finding in the context of the evolutionary ‘duplication and migration’ hypothesis (Belyk and Brown, 2017; Jarvis, 2019) and conclude that their finding suggest a primary role of the dLMC for laryngeal motor control in primary motor cortex. Another study determines the relationship between structural connectivity, cortical myelin content, and cortical thickness (Bajada *et al*., 2019). Cortical areas that assemble short range fibers have relatively high myelin content and lower cortical thickness whereas cortical areas with long range fibers have relatively low myelin and a higher cortical thickness. Furthermore, numbers of short range fibers are proportionally greater in primary cortical areas, whereas numbers of long range fibers are proportionally greater in associated cortical areas. The authors suggest that cortical thickness may vary with fiber length because a thicker cortex may allow more fibers to converge. Although, diffusion-weighted tractography is widely accepted as a valid method to assess white matter connectivity *in vivo* in humans (Haber *et al*., 2020), it is important to keep in mind that various caveats bias tractography data (Van Essen *et al*., 2014) and validation by invasive studies is desirable. Still, in light of the findings from cortical myelin mapping of the laryngeal representations (Eichert *et al*., 2020) and fiber length profiling (Bajada *et al*., 2019), our finding of diverging connectivity profiles of the two laryngeal representations with the dLMC to show a denser structural network compared to the vLMC seems plausible.

Differential patterns of structural connectivity of the two laryngeal representations also include hemispheric lateralization. Both LMCs demonstrated right lateralization of the superior temporal gyrus, and left lateralization of the caudate nucleus, which is consistent with the directions reported in the previous report on dLMC connectivity (Kumar *et al*., 2016). In addition, while the dLMC demonstrated left lateralization of the anterior portion of the inferior frontal gyrus pars opercularis, also consistent with the previous report, the somatosensory cortex, the posterior portion of the inferior frontal gyrus pars opercularis and the somatosensory cortex were right lateralized for vLMC. This heterogeneity also with respect to a hemispheric specialization supports the idea of a functional dissociation of the two laryngeal representations (Belyk and Brown, 2017; Dichter *et al*., 2018; Eichert *et al*., 2020). In particular, the modulation of pitch in speech and singing has been suggested to be primarily controlled via the dorsal laryngeal motor cortex (Dichter *et al*., 2018; Eichert *et al*., 2020). And causal inference with transcranial magnetic stimulation demonstrated, for example, that in particular the laryngeal representation in the right hemisphere is involved in vocal pitch regulation (Finkel *et al*., 2019) and auditory pitch discrimination (Sammler *et al*., 2015). However, currently the ground truth of the anatomical connectivity of LMC to laryngeal motor neurons and cortical and subcortical brain areas results from tracing studies of a single cortical motor representation in mammals and humans (Kuypers, 1958; Kirzinger and Jürgens, 1982; Simonyan and Jürgens, 2002; Simonyan, 2014). Thus, neurophysiological and brain stimulation studies (Hamdy *et al*., 1998) might be advantageous to map out the particular connectivity of a dual representation in humans, and to foster the distinct roles of dLMC and vLMC in concordant and specific larynx functions.

A functional dissociation of the laryngeal representations is further suggested by the varying involvement of these two areas in persistent developmental stuttering. This speech fluency disorder is characterized by a white matter deficit, i.e. a reduced fractional anisotropy, in the left vLMC (Sommer *et al*., 2002; Chang *et al*., 2008; Watkins *et al*., 2008) and most likely involves fibers of the superior longitudinal fasciculus/arcuate fasciculus (Connally *et al*., 2014; Neef *et al*., 2015, 2018; Kronfeld-Duenias *et al*., 2016, 2018). The white matter deficit might cause a disconnection of the ventral laryngeal motor representation from left perisylvian speech regions (Sommer *et al*., 2002). Here, we substantiated this longstanding finding by showing that adults who stutter have a reduced overall connection probability of the vLMCs when compared to fluent speakers. Moreover, structural connectivity profiling of both laryngeal motor representations revealed that only vLMC structural connectivity serves as a powerful statistical predictor of stuttering severity. In particular, connection probability of the left vLMC with the left primary somatosensory cortex and inferior gyrus pars opercularis, and connectivity of the right vLMC with the right primary somatosensory cortex, inferior parietal lobe, putamen and caudate nucleus strongly related to the motor signs of stuttering. The involvement of frontal and parietal sites substantiates the assumption that stuttering results from an insufficient feedforward and feedback control during speech related sensorimotor signal transmission (Guenther, 2016). In addition, our findings further affirm that affected circuits extend beyond the known left hemisphere speech motor pathways (Kronfeld-Duenias *et al*., 2018; Neef *et al*., 2018) and engage the basal ganglia system (Alm, 2004; Connally *et al*., 2018; Chang and Guenther, 2019).

Both LMC networks established strong connections with cortical brain areas specified to process planning and timing of motor sequences, sensory input and feedback, and sensorimotor integration. Strikingly, only the ventral laryngeal representation is affected in stuttering. Our vLMC seed coordinate was derived from the first dMRI study on stuttering [x = -48, y = -15, z = 18] (Sommer *et al*., 2002). This white matter site is closely located to sites of cortical activity reported for tasks that were designed to stimulate and differentiate dLMC and vLMC brain activity during whistling and singing [x = -59, y = -16, z = 13] (Belyk *et al*., 2020) or vocalization and vowel production [x = -58, y = -2, z = 20] (Eichert *et al*., 2020). Likewise, earlier fMRI studies that investigated vowel production (Grabski *et al*., 2012), vocal imitation (Belyk *et al*., 2016), pitch (Peck *et al*., 2009), cough (Mazzone *et al*., 2011), brain alterations in spasmodic dysphonia (Simonyan and Ludlow, 2012) relate laryngeal control to our chosen vLMC coordinate. Cyto- and myeloarchitecture of the ventral laryngeal representation is currently unknown and it has been suggested that this region might belong to the cytoarchitectonic area 6 (Pfenning *et al*., 2014; Dichter *et al*., 2018; Jarvis, 2019; Eichert *et al*., 2020) or to the cytoarchitectonic area 43 (Belyk and Brown, 2017; Belyk *et al*., 2020). Besides this dissent, different research groups seem to agree on the idea that the vLMC does not belong to the primary motor cortex. The assignment of the two laryngeal representations to different architectural areas underpins the suggestion of segregated functions. However, it remains unclear how the vLMC contributes distinctively to larynx control.

One striking phenomenon in stuttering is the preserved ability to sing. In contrast, speech prosody and pitch control are apparently disrupted during stuttering (Neumann *et al*., 2018). Both vocal functions, singing and speaking, involve a dedicated control of laryngeal muscles to regulate pitch and voicing and to coordinate vocalization with articulation and breathing. Likewise, both functions rely on shared cognitive processes and large-scale networks that overlap to a great extend (Özdemir *et al*., 2006). However, a theoretical discussion infers that musical pitch requires a more accurate encoding to ensure discrete melody production than does speech, for which pitch variation is continuous (Zatorre and Baum, 2012). Accordingly, less degrees of freedom for pitch modulation in singing provide a finer and more specified template of upcoming vocalizations. Such a temporal specification that also includes rhythm in song, might facilitate fluency as observed when affected individuals sing. The same reasoning holds true for other fluency enhancing techniques such as chorus reading and metronome speaking (Barber, 1940; Wingate, 1969; Davidow *et al*., 2009), carry-over fluency induced by extreme prolongations (Briley *et al*., 2016), or reduced voicing complexity by fluency-shaping (Euler *et al*., 2009). It seems plausible to assume that network formation for speech and song production is dynamic and task-dependent and varies in concert with involved brain regions that perform parallel computations. Singing might recruit network formations with a widely intact structural organization biased by a stronger dLMC involvement, while speaking might more heavily involve vLMC networks, which are affected in stuttering. A different interpretation can be drawn from a recent case study. Two expert musicians underwent awake craniotomy surgery. The stimulation of the vLMC area disrupted speech and music production, i.e. playing the piano or the guitar (Leonard *et al*., 2019). The authors suggest that this ventral area might code more complex representations that are independent of specific effectors such as laryngeal muscles.

The current analysis revealed no impact of an intensive fluency-shaping intervention on white matter networks of the two laryngeal representations. This result is somewhat counterintuitive because the speech restructuring method required individuals who stutter to learn a changed speech pattern. This speech pattern comprised soft voice onsets, consonant lenitions, and controlled sound prolongations. Thus, voicing and timing were the key features under change over the course of the acquisition of the new speech technique (Euler *et al*., 2009). In contrast to unchanged white matter structures of voice control, resting-state connectivity was strengthened within the dLMC network (Korzeczek *et al*., 2020). Specifically, intervention synchronized resting-state activity between the left dLMC and the left posterior portion of the inferior frontal gyrus pars opercularis, the left inferior parietal lobe, and the right posterior superior temporal gyrus. The observation that the structural dLMC network is unaffected in stuttering, but recruited by stuttering intervention suggest a compensatory involvement of this networks in assisted recovery. This is a new finding that contradicts previous reports on an intervention-induced normalization of brain activity (Neumann *et al*., 2005, 2018; Kell *et al*., 2009, 2017).

In sum, present findings strongly support the view of a functional segregation of the dual cortical larynx representations, which is based on a diverging structural network organization. The dorsal laryngeal representation has an overall denser structural network compared to the ventral one. The intrahemispheric connectivity profiles of bilateral ventral laryngeal representations predict motor signs of stuttering over and above the biological variates age, sex, and handedness, and serves as a weighty neuronal trait marker of stuttering. However, the vLMC network is insensitive to intensive fluency-shaping, i.e. shows no structural neuroplasticity after restructured pitch and voicing in speech.

## Abbreviations

aIFGop: anterior inferior frontal gyrus pars opercularis
ANCOVA: analysis of covariance
aSTG: anterior superior temporal gyrus
Caud: caudate nucleus
dLMC: dorsal laryngeal motor cortex
Gp: globus pallidus
IPL: inferior parietal lobe
pIFGop: posterior inferior frontal gyrus pars opercularis
Put: putamen
S1: primary somatosensory cortex
SMA: supplementary motor area
vLMC: ventral laryngeal motor cortex

## 5. Acknowledgments

We thank Bettina Helten for the co-analysis of the speech samples, Michael Bartl for supporting the organization of the behavioural data, and Britta Perl and Ilona Pfahlert for assistance with the acquisition of the MRI data.

## Author’s Roles

NEN, AP, WP and MS research project conception; NEN data analyses, statistical analyses design and execution, writing of the first draft; AP data acquisition; AP, AWvG, PD, HCR, WP and MS analyses review and critique, manuscript review and critique; MS and WP overall study coordination and funding.

